# Autonomous development and regeneration of isolated rice egg cells in a fertilization-independent manner

**DOI:** 10.1101/2023.05.02.539102

**Authors:** Kasidit Rattanawong, Kaori Totsuka, Shizuka Koshimizu, Kentaro Yano, Takashi Okamoto

## Abstract

Parthenogenesis is suppressed in rice egg cells to avoid precocious development before fertilization. We found that cold treatment released cell cycle arrest in egg cells and triggered mitosis. Egg cells isolated from *japonica* (Nipponbare; NB) and *aus* (Kasalath; KS) subspecies divided and regenerated into mature plants after cold treatment. The egg-derived plants showed variety of ploidy levels, including haploid (n), diploid (2n), and tetraploid (4n). Nuclear DNA quantification showed that genome duplication occurred during early parthenogenetic development. Owing to the presence of single nucleotide polymorphisms (SNPs) between NB and KS, inter-subspecific hybrid plants (NB-KS hybrids) were created via electrofusion. Egg cells from the NB-KS hybrid developed parthenogenetically into polyploid plants. 2n and 4n plants originating from the same NB-KS egg cell displayed the same homozygous SNP patterns throughout the genome, indicating that these plants were doubled and quadrupled haploids. Transcriptome analyses of cold-treated egg cells demonstrated that parthenogenesis-related candidate genes, including *OsBBML1*, were upregulated.

## Introduction

Living organisms generally expand the range of their species via sexual reproduction, which requires the contribution of both male and female gametes. The resulting higher levels of genetic diversity lead to an increase in fitness and long-term survival of a species through the combination of allele sets from both parents known as “biparental inheritance.” The production of genetically identical clones through asexual reproduction is crucial for a variety of species that require rapid and substantial increases in the number of offspring. Because only a single parent is required for asexual reproduction, the created progeny will possess full sets of genes derived from one parent known as “uniparental inheritance.” Asexual reproduction can proceed via somatic or gametic cells. A well-studied female gamete-mediated embryogenesis that widely occurs in plant and animal species is parthenogenesis, spontaneous development of an embryo from an unfertilized egg cell (reviewed in Vijverberg, Ozias-Akins, and Schranz 2019). Although the emergence of progenies from unfertilized oocytes or egg cells is a common phenomenon in animals and plants, the production mode of parthenogenetic progenies differs between animals and flowering plants. In animals, autonomous development of the parthenogenetic embryo occurs after meiosis with or without restoration of diploid chromosome number via fusion of haploid nuclei or chromosome doubling during the endomitotic cycle (Avise, 2008). The nutrient supply during embryonic development is directly provided by the mother, resulting in the formation of mature embryos as parthenogenetic progeny. In flowering plants, progenies are formed as seeds consisting of embryos and endosperms, which are derived from fertilized egg cells and central cells; hence, parthenogenetic development of unfertilized egg cells can be detected as a part of fertilization-independent asexual seed formation known as apomixis. In diplospory, an apomictic type of unreduced female gametes are produced via apomeiosis, and unreduced egg cells parthenogenetically develop into embryos with autonomous or pseudogamous development of unreduced central cells into the endosperm (Pupilli and Barcaccia, 2012; Conner et al., 2017; Hand and Koltunow, 2014).

Investigations of parthenogenesis in angiosperms have been conducted mainly using three approaches: 1) cytological analyses, 2) genetic approaches, and 3) autonomous development of egg cells. Cytological studies have indicated that in angiosperms, mature egg cells normally possess largely condensed repressive chromatin and a relatively silent transcriptional state, a mechanism that prohibits egg cells from precocious development without proper fertilization with a sperm cell (Garcia-Aguilar et al., 2010; Pillot et al., 2010). These repression mechanisms need to be released to enable developmentally arrested egg cells to proceed with parthenogenesis or even undergo normal zygotic development to obtain totipotency after fertilization (Baroux and Grossniklaus, 2015). It is assumed that parthenogenesis is triggered by genetic reprogramming via spontaneous de-repression of chromatin and subsequent activation of transcription machinery in unfertilized egg cells. In addition, cell cycle arrest is abrogated or highly reduced in parthenogenetic egg cells.

Genetic approaches to study parthenogenesis have been based on the determination of parthenogenesis-related gene loci and subsequent identification of genes responsible for parthenogenesis in the loci. Genes encoding transcription factors *Apospory-specific Genome Region BabyBoom*-like (*ASGR-BBML*) 1 and *PARTHENOGENESIS* (*PAR*) were identified in parthenogenetic pearl millet (*Pennisetum squamulatum*) and dandelion, respectively, and have been reported to induce parthenogenesis in egg cells of sexual pearl millet and dandelion upon ectopic expression under *Arabidopsis* egg cell-specific promoters (Conner et al., 2015; Underwood et al., 2022). In addition, *PsASGR-BBML1* and *OsASGR-BBML1* (*OsBBML1*), rice orthologs of *PsASGR-BBML1*, induce parthenogenesis in rice and maize through their ectopic expression in egg cells (Conner et al., 2017; Khanday et al., 2019; Rahman et al., 2019). Heterologous expression of *PAR* gene from the dandelion, driven by *EC1* promoter, induced the formation of parthenogenetic haploid embryos in lettuce egg cells without fertilization (Underwood et al., 2022).

The developmental profile of parthenogenetic egg cells was monitored with the Salmon System established in wheat, as the “Salmon” line, in which the short arm of chromosome 1B of wheat is replaced by the short arm of chromosome 1R of rye, produces haploid embryos via parthenogenesis where the cytoplasm of *Aegilops kotschyi* or *Ae. caudata* is introduced (Tsunewaki and Mukai, 1990). It has been demonstrated that parthenogenetic development of egg cells is independent of the signals derived from ovular tissues/ovaries, as embryos could develop directly from egg cells isolated from parthenogenetic lines of Salmon wheat, (*caudata*)-Salmon, and (*kotschyi*)-Salmon, in an in vitro culture system (Tsunewaki and Mukai, 1990; Kumlehn et al., 2001). Transcriptional analysis using a cDNA library synthesized from mRNA isolated from parthenogenetic egg cells of caudata-Salmon wheat showed that the highly represented ETS group was a homologue of the barley ECA1 gene, which is specifically expressed in the early stage of microspore embryogenesis, suggesting its potential role in the parthenogenesis of Salmon wheat (Kumlehn et al., 2001; Pulido et al., 2009).

Within embryo sacs, plant egg cells are widely known to reside in quiescence at the G1 phase of the cell cycle but appear to be ready for development upon fertilization in both monocots and dicots (Mogensen and Holm, 1995; Sukawa and Okamoto, 2018; Liu et al., 2020). Notably, single-cell type transcriptome analyses using egg cells of *Arabidopsis* (Wuest et al., 2010), rice (Anderson et al., 2013), and maize (Chen et al., 2017) suggested that RNA, proteins, and other molecules are stored in egg cells to support embryogenesis upon egg activation. Therefore, a single factor might be sufficient to release cell cycle arrest and activate embryogenic development, thereby initiating the parthenogenetic (autonomous) development of egg cells (Vijverberg et al., 2019). Initiation of parthenogenesis induced by the ectopic expression of *ASGR-BBML1* and *PAR1* in egg cells is an artificial parthenogenetic trigger (Khanday et al., 2019; Rahman et al., 2019; Underwood et al., 2022).

As possible triggers for breaking the repressive state of egg cells, environmental stresses and stimuli have been reported as parthenogenesis-initiating factors. For example, ovaries isolated from sexual plants such as durum wheat (Sibi et al., 2001) and sugar beets (Gürel et al., 2000) were pretreated with cold stress and then cultured to obtain haploid plants. Although direct evidence is still lacking, it has been suggested that this phenomenon, termed gynogenesis, is due to the parthenogenetic development of egg cells embedded in cold-stressed ovaries. Parthenogenesis in angiosperms has been intensively investigated, as asexual events are closely related to the nature of plant reproduction and are highly important for agricultural use in the production of doubled-haploid plants for instant genotype fixation (heterosis fixation) (Sailer et al., 2016). However, knowledge of the mechanisms of parthenogenesis in angiosperms remains limited. This is partly because the egg cells of angiosperms are deeply embedded in the ovaries, making direct analysis of the egg cell difficult. Moreover, an experimental system that can artificially induce autonomous division and development of egg cells has not yet been established.

In this study, we successfully demonstrated the autonomous division and development of isolated rice egg cells from unpollinated ovaries in a fertilization-independent manner. Despite residing in quiescence, the exposure of egg cells isolated from wild-type *japonica* rice (*Oryza sativa* cv. Nipponbare, NB) to cold stress can effectively induce spontaneous division. In addition, we found that egg cells isolated from *aus* rice (*O. sativa* cv. Kasalath, KS) showed parthenogenetic development without cold-pretreatment. The divided egg cells were able to regenerate into sexually mature plants with a variety of ploidy levels, including haploid (n), diploid (2n), and tetraploid (4n), with variable morphological characteristics and fertility. Nuclear DNA quantification was performed to demonstrate that genome duplication from n to 2n occurs during the early egg cell proliferation stage. Inter-subspecific hybrid plants between NB and KS (NB-KS hybrid) were created via electrofusion of NB egg and KS sperm cells. The egg cells isolated from the NB-KS hybrid plants were cultured into plantlets, resulting in the production of haploid, diploid, and tetraploid rice plants. Genome sequencing and subsequent single nucleotide polymorphism (SNP)-based analyses of these rice plants confirmed that diploid and tetraploid plants from isolated rice egg cells are doubled and quadrupled haploids, respectively. Finally, transcriptome analyses of isolated egg cells treated with cold temperature revealed upregulation of parthenogenesis-inducing genes and downregulation of dormancy-related genes, indicating the possible conversion of cold-treated egg cells from quiescent to parthenogenetic states.

## Results

### Parthenogenetic development of egg cells isolated from unpollinated flowers of *japonica* (cv NB) and *aus* (cv KS) rice plants

To examine the parthenogenetic competence of sexual non-parthenogenetic *japonica* rice (cv NB), egg cells, which reproduce through sexual pathways and require fertilization with sperm cells for activation toward embryogenesis, were isolated from unpollinated flowers and cultured to examine the possible fertilization-independent division and development of egg cells. When isolated NB egg cells were transferred into the culture medium without any pretreatment, all 30 cultured egg cells became undeveloped and degenerated within three days after culture (DAC) (Fig. 1 and Table 1). However, when egg cells were pre-incubated at 4 °C for 12 h prior to cultivation, 25 of 216 cultured cells (11.5%) divided into 2-celled structures at 2 DAC and developed further into multicellular structures within 5 DAC (Figs. 1, 2, and Table 1). These results suggest that preceding cold treatment can trigger the parthenogenetic development of isolated rice egg cells without the contribution of male factors and independent of the signals derived from ovary/ovular tissues. Notably, 19 cell masses were developed from the 25 autonomously divided egg cells; and among these, 16 further developed into white cell colonies (Fig. 2 and Table 1), which were then transferred to regeneration medium to stimulate shoot and root formation. Eight regenerated plantlets were obtained (Fig. 2 and Table 1), which were then transferred to soil pods and cultivated in an environmental chamber, allowing them to grow into mature rice plants with developed flowers (Supplementary Fig. 2).

**Figure 1.**
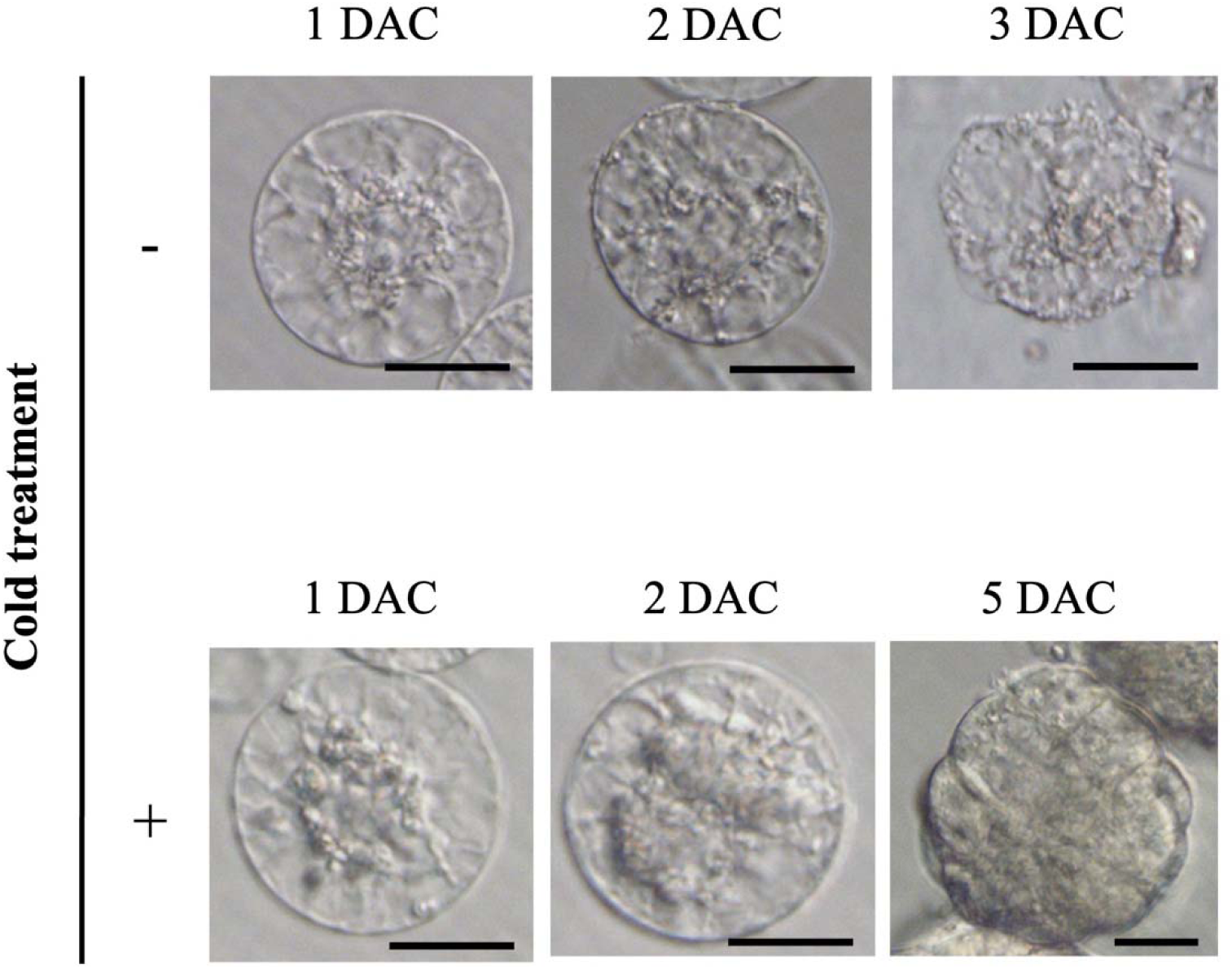
Autonomous cell division profile of isolated NB egg cells cultured with or without preceding 12-h cold treatment. Minus (−) and plus (+) signs indicate egg cells cultured without or with preceding cold treatment for 12 h, respectively. Scale bars: 20 μm.

**Figure 2.**
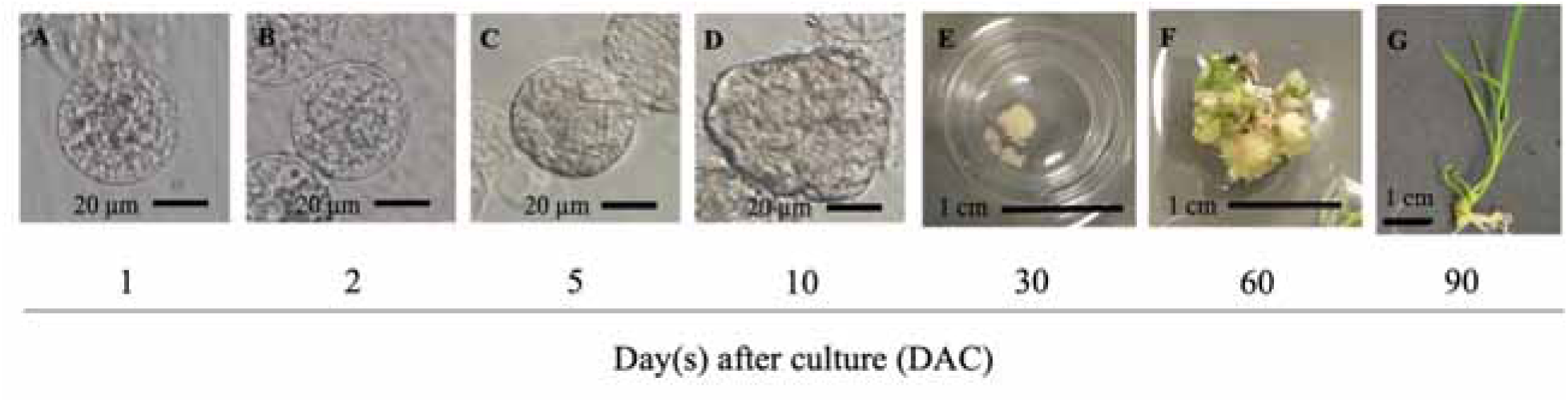
Development and regeneration plantlets developed from isolated NB egg cells exposed to 12 h-cold treatment before cultivation in growth and regeneration media. (**A**) Cold-treated egg cells remained in a single-celled stage one day after culture (DAC). (**B**) Nuclear/cell division was observed at two DAC. (**C**, **D**) Divided egg cells developed into multicellular structures at about 5 DAC. (**E**) White cell colonies were formed within 30 DAC, and (**F**) leafy shoots and roots were regenerated upon transfer to the regeneration medium at 60 DAC. (**G**). Mature plantlets were subsequently obtained at about 90 DAC, which were then cultivated in a soil pod.

**Table 1.**
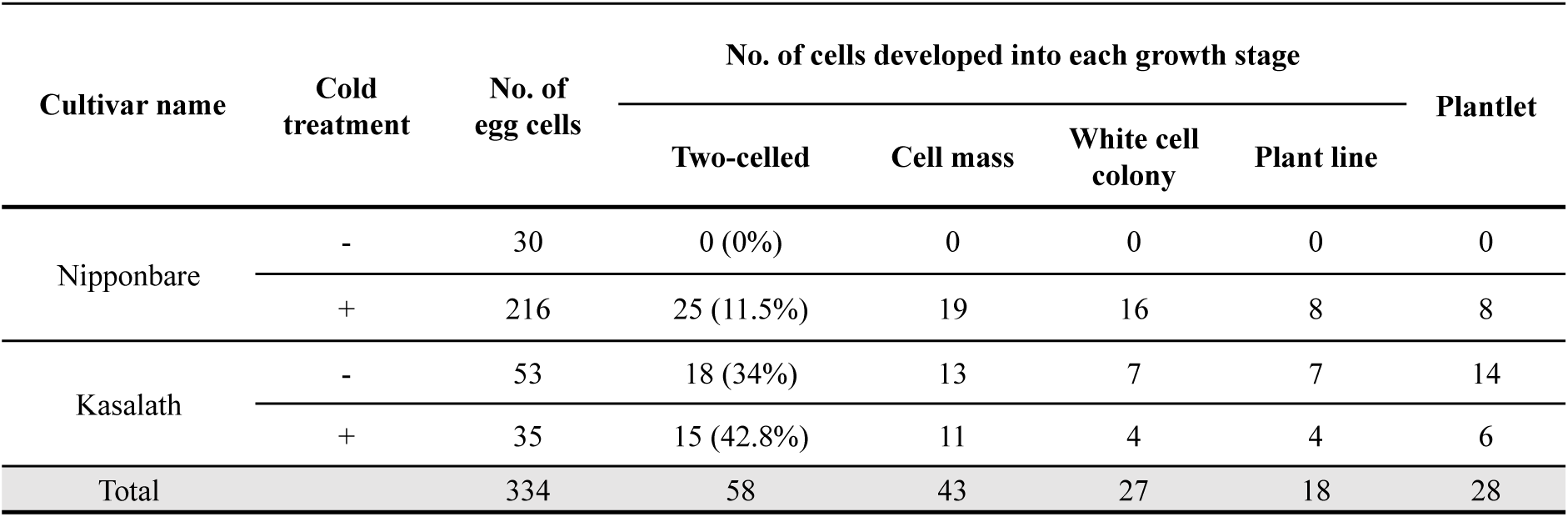
Developmental profiles of Nipponebare and Kasalath egg cells cultured with or without preceding cold treatment prior to cultivation.

In addition to *japonica* NB rice, we tested the parthenogenetic competence of egg cells isolated from unpollinated flowers of *aus* rice (cv. KS) using the same experimental procedure. Interestingly, 18 of 53 KS egg cells (34%) exhibited cell division without cold treatment (Table 1), and in the case of KS egg cells with cold treatment, the division rate increased to 42.8% (Table 1). This suggests that regardless of cold exposure, egg cells isolated from unpollinated flowers of KS rice plants showed high efficiency of fertilization-independent development and that KS egg cells might be more parthenogenetically active than NB egg cells under the in vitro culture system. Moreover, it was also suggested that the procedure of egg cell isolation from unpollinated ovaries and subsequent culture of isolated cells can trigger parthenogenetic development of KS egg cells, and that pretreatment with cold stress further enhances their parthenogenetic potential. To overcome the difficulty in visualizing the number of nuclei/cells during autonomous egg cell division, KS plants expressing GFP-tagged histones (H2B-GFP) were produced to enable direct observation of the number of nuclei during the parthenogenetic development of egg cells (Abiko et al., 2013). We found that without cold treatment egg cells started to divide at 2 DAC, as two nuclei were detected in the divided egg cells and multiple nuclei of the multicellular structures were observed at 3–4 DAC (Supplementary Fig. 1A). A single nucleus was continuously visualized in the undivided egg cells, and the cells disintegrated at about 4 DAC (Supplementary Fig. 1A). These results suggest that the first cell division of parthenogenetically activated egg cells appeared at 1 DAC. For the number of plants formed from divided egg cells, without cold treatment, 13 cell masses developed into seven white-cell colonies, which later formed seven plant lines (Table 1). Because a plant line has the potential to regenerate into multiple shoots under the callus-mediated regeneration system used in this study (Uchiumi et al., 2007; Toki et al., 2006), each KS egg-derived plant line regenerated into two plantlets, resulting in the formation of 14 plantlets (Table 1). For the cold-treated KS egg cells, 4 of 11 cell masses developed into white cell colonies, and four plant lines were obtained, which were regenerated into six plantlets (Table 1).

### Characterization of egg cell-derived rice plants: Ploidy, morphological characteristics, fertility, and gamete isolation efficiency

The ploidy levels of the rice plants regenerated from egg cells were measured to examine whether the original ploidy level of egg cells was retained. First, nuclei were extracted from the leaves of diploid wild-type NB plants; the DNA content per nucleus was measured via flow cytometry, which normally exhibited one single peak of 2C, and used as a diploid control for ploidy level assessment (Fig. 3A). Of the 28 egg-derived plants (Table 1), four plants with small flowers and short statures (Fig. 3H, I, J, Table 2, and Supplementary Fig. 2) showed a peak at 1C (Fig. 3B), whereas 10 and 12 plants exhibited 2C and 4C peaks, respectively (Fig. 3C, D and Table 2). Peaks beyond 4C were also detected in the two egg-derived plants, which were estimated to be aneuploid (Fig. 3F and Table 2). These results suggest that the plants regenerated from egg cells preferably undergo genomic duplication to increase the ploidy numbers, as three-fourths of the plants obtained from egg development were shown to possess ploidy levels higher than haploids. In addition, the ratio of 2n:4n was almost equal to 1:1 (Table 2), indicating that repeated genome duplication occurred during the culture of isolated egg cells and/or regeneration of egg cell-derived cell masses.

**Figure 3.**
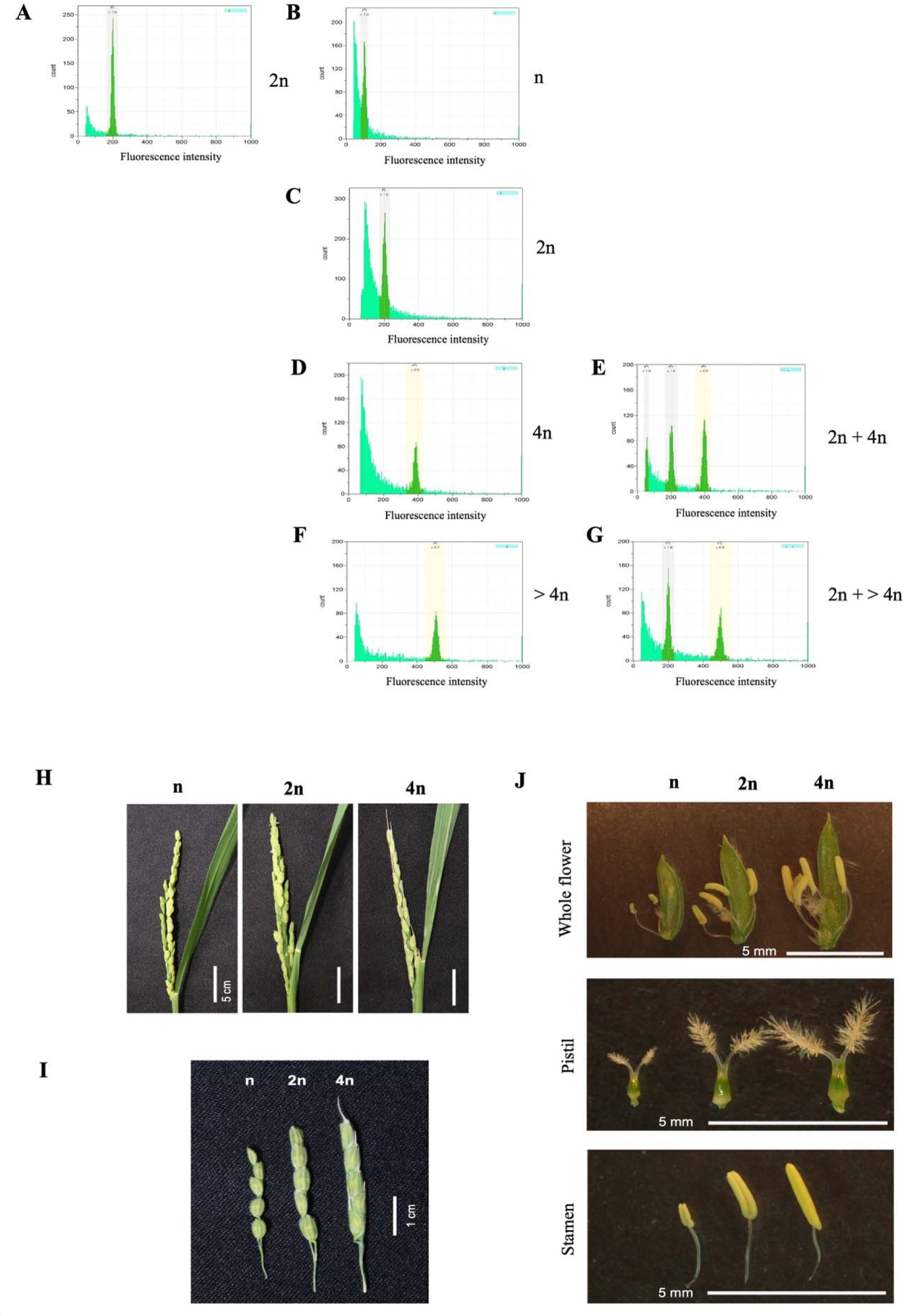
Ploidy levels and morphological characterization of reproductive structures of egg cell-derived plants. (**A**) Nuclei extracted from the leaves of wild-type rice plants and (**B**–**G**) plants regenerated from NB egg cells subjected to 12-h cold temperature pretreatment; the DNA content per nucleus was measured via flow cytometry. **B**, **C**, and **D** indicate the haploid, diploid, and tetraploid amounts of DNA, respectively. F indicates the amount of DNA beyond 4n, which was identified as aneuploidy. (**E**, **G**) Nuclei extracted from wild-type rice plants were loaded as an internal control, along with those from (**E**) tetraploid and (**G**) aneuploid plants. (**H**, **I**) Comparison of morphological features of (**H**) panicles and (**I**) florets of haploid (n), diploid (2n), and tetraploid (4n) rice plants regenerated from cold-treated NB egg cells. (**J**) Flowers from egg cell-derived plants were dissected to observe differences in morphological features of reproductive structures: whole flowers (top panel), pistils (middle panel), and stamens (bottom panel) isolated from haploid (n), diploid (2n), and tetraploid (4n) plants.

**Table 2.**
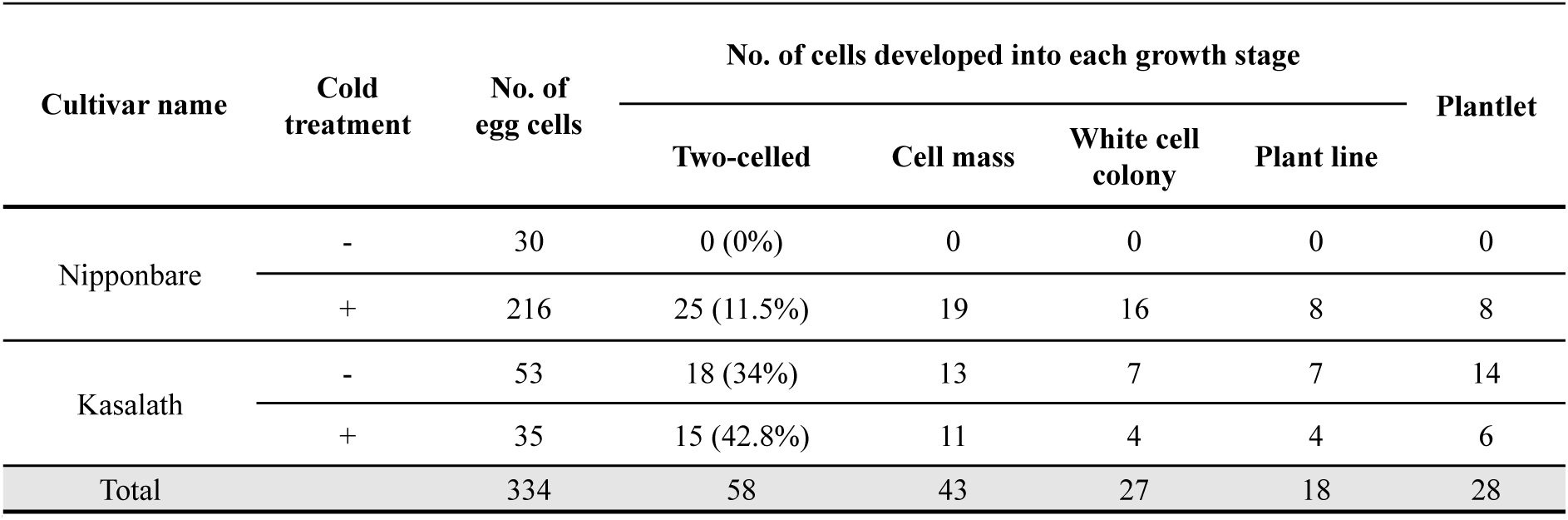
Ploidy levels of plants derived from egg cells isolated from Nipponbare and Kasalath plants.

### Genome duplication during early proliferation of NB-derived egg cells treated with or without cold treatment

The presence of haploid individuals confirms that the egg-derived plants were directly developed from the haploid egg cells with cold treatment of NB egg cells and with or without cold treatment of KS egg cells (Fig. 2 and Table 2). Notably, the majority of egg-derived plants were diploid and tetraploid (Table 2), suggesting that genome doubling might occur during the parthenogenetic development of egg cells into plantlets. Therefore, to judge whether the possible genome duplication occurred during the early proliferation stage of the egg cell, the relative amount of nuclear DNA in egg cells and the cells of 2-celled, 4- to 8-celled, and 10- to 20-celled stages of egg-derived multicellular structures was monitored (Fig. 4). The frequency histogram of the relative amount of nuclear DNA of the haploid egg cells showed fluorescence intensity grouped around 3 arbitrary units (AU), which corresponds to the haploid amounts of nuclear DNA (Fig. 4A). In the 2-celled stage, the frequency histogram indicated the emergence of cells that exhibited an approximately 2-fold increase in relative fluorescence intensity compared to the egg cells (Fig. 4B). The presence of a larger number of cells with increased relative fluorescence intensity was observed in 4- to 8-celled multicellular structures (Fig. 4C), and the increased relative fluorescence intensity was detected in the nuclei of most cells of 10- to 20-celled multicellular structures (Fig. 4D). The gradual increase in the percentage of cells possessing approximately double the amount of nuclear DNA compared to egg cells suggested that duplication of the nuclear genome occurred during fertilization-independent development of egg cells into multicellular structures (Fig. 4A–D). Taken together, during the early stage of parthenogenetic development, the genome of haploid egg cells doubled, and the ploidy level of developing egg cells increased 2-fold, thereby changing from a haploid to diploid state.

**Figure 4.**
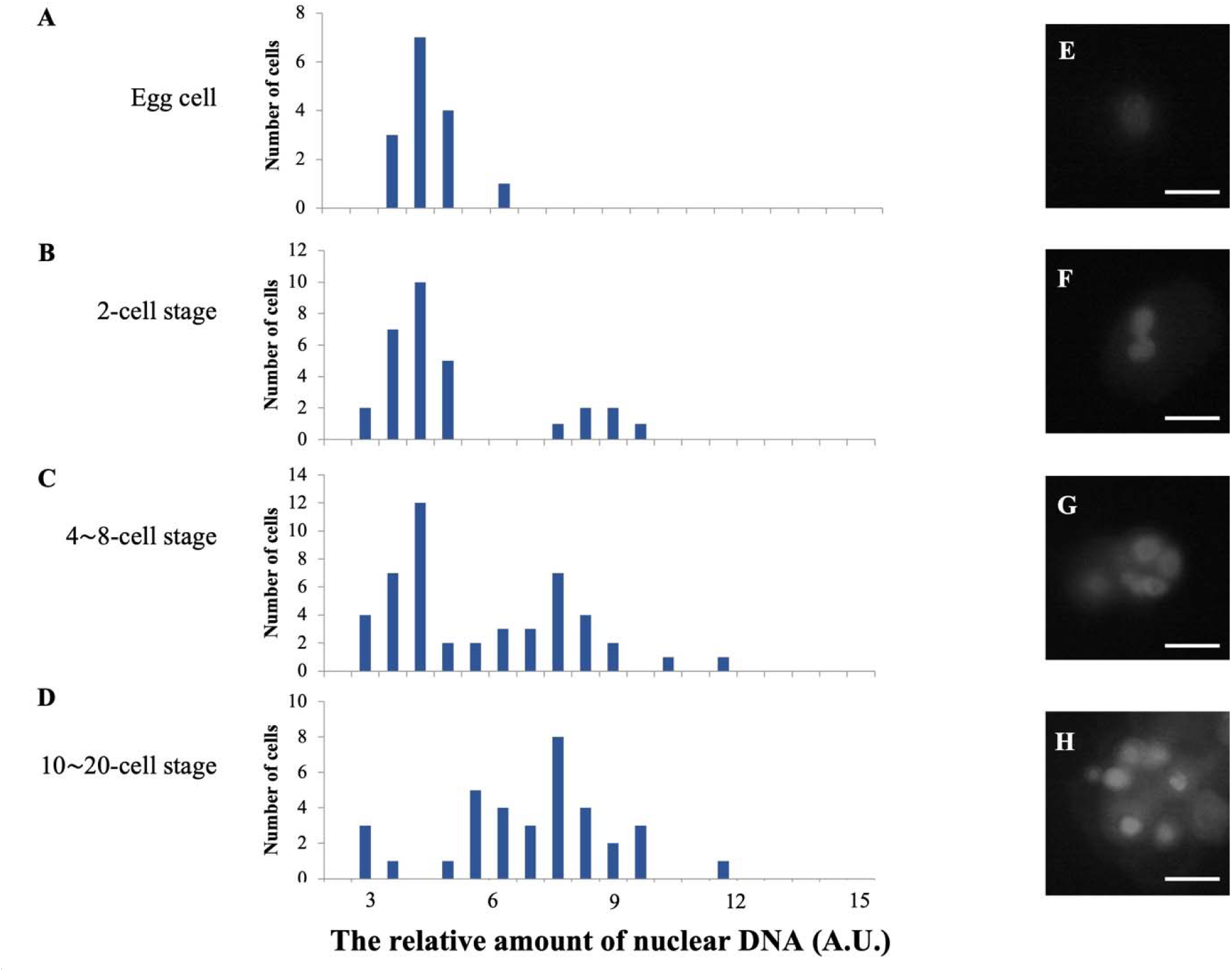
Estimation of ploidy levels through direct visualization of nuclear DNA contents dividing egg cells and developing multicellular structures. Modified DAPI staining of nuclear DNA of egg cells and the cells of 2-celled, 4- to 8-celled, and 10- to 20-celled stages of egg-derived multicellular structures (Sukawa and Okamoto 2018). (**A**–**D**) Fluorescence intensity derived from the (**A**) egg, (**B**) 2-celled, (**C**) 4- to 8-celled, and (**D**) 10-to 20-celled stages was quantified and plotted as histograms. Fluorescence images showing fluorescence intensity representing the amount of nuclear DNA in (**E**) egg cells, (**F**) 2-celled, (**G**) 4- to 8-celled, and (**H**) and 10- to 20-celled stages. Scale bars in **E**-**H**: 20 μm.

### Production of NB-KS hybrid rice plants, culture of egg cells isolated from NB-KS hybrid, and ploidy of rice plants regenerated from the NB-KS hybrid egg cells

By culturing NB egg cells with cold treatment and KS egg cells, haploid, diploid, and tetraploid rice plants were obtained, and possible genome polyploidization was considered to occur during egg cell proliferation (Fig. 4). Further, to confirm that these polyploids are true doubled-haploid and quadrupled-haploid at the genomic DNA level, we prepared hybrid rice plants between NB and KS. Egg cells (NBKS egg cells) isolated from the unpollinated flowers of the NBKS hybrid rice plants were cultured into plants, and the genome DNA sequences from these regenerated plants were analyzed using SNPs between NB and KS. An NB egg cell was fused with a KS sperm cell, and the resultant NB-KS hybrid zygote was cultured into plantlets, termed the NBKS hybrid. Three NBKS hybrid lines derived from three independent NBKS zygotes, termed NBKS hybrids I, II, and III, were used for the developmental profile investigation. When egg cells isolated from the NBKS hybrid, termed NBKS egg cells, were cultured with or without cold pretreatment, they parthenogenetically developed and regenerated into plantlets (Fig. 5), with a division efficiency of 14–26% depending on the treatment conditions (Table 3). The ploidy of all regenerated plants (126 plants, Table 4) was measured, and 10 haploid, 51 diploid, and 54 tetraploid plants were detected (Table 5). To examine the doubling of haploid embryos at the genome sequence level, several regenerated plants were subjected to genome sequence analysis.

**Figure 5.**
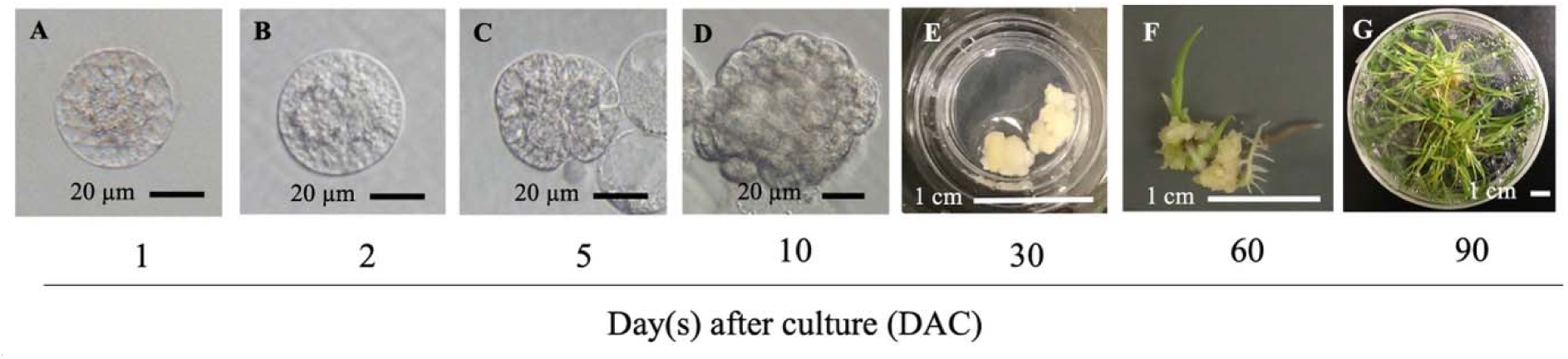
Development and regeneration of plantlets derived from egg cells isolated from NBKS hybrid plants. (**A**) Egg cells isolated from NBKS plants were treated with or without cold temperature for 12 h prior to culturing in the growth and regeneration media. The cultured egg cells remained in a single-celled stage one day after culture (DAC). (**B**) Cell/nuclear division was then observed at 2 DAC, and (**C**, **D**) the formation of multicellular structures was visible from 5 DAC onwards. (**E**). Developing cell mass formed white-cell colonies at about 30 DAC. (**F**) Leafy shoots and branched roots regenerated at about 60 DAC. (**G**) Mature plantlets were formed and transferred to a soil pod.

**Table 3.**
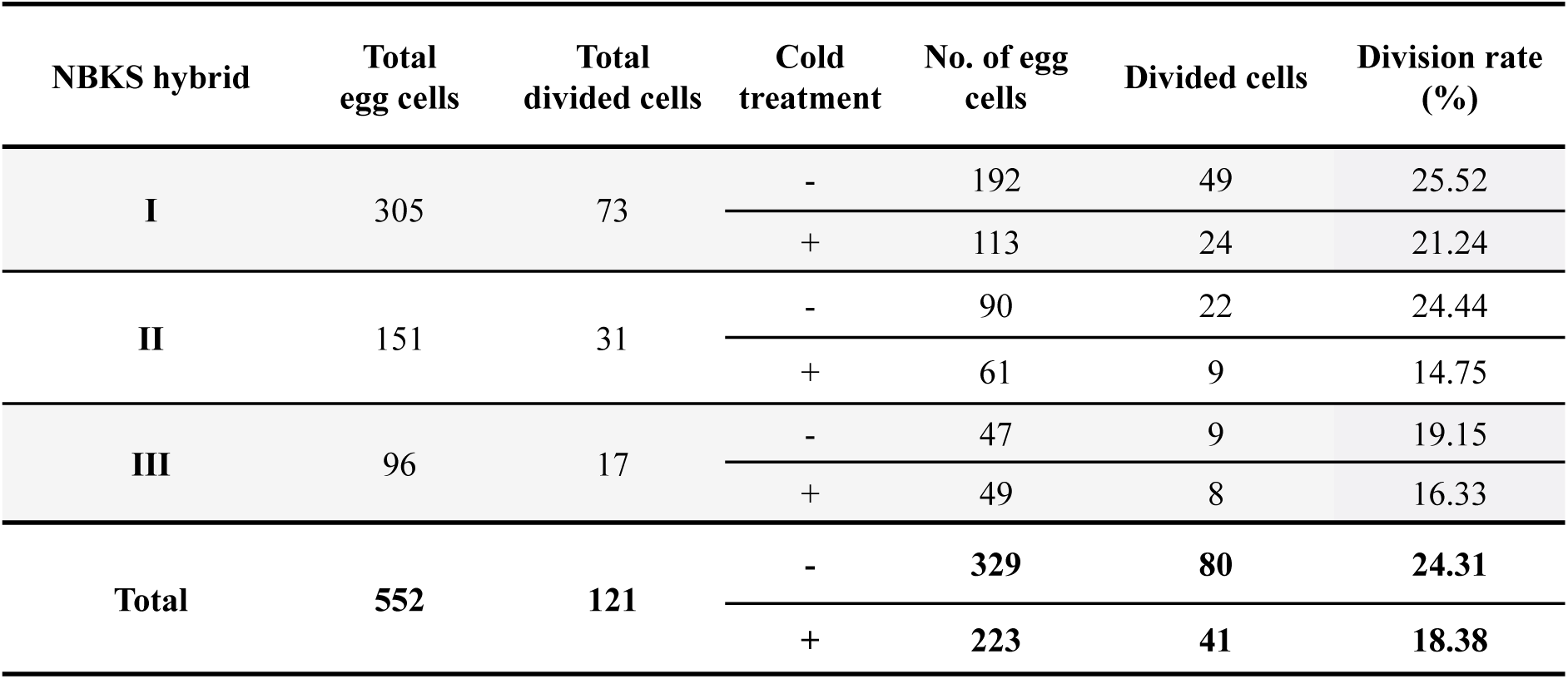
Rates of autonomous cell division of egg cells isolated from NBKS hybrid plants cultured with or without preceding cold treatment.

**Table 4.** Developmental profiles of egg cells isolated from NBKS plants cultured with or without 12-h cold treatment.

| NBKS hybrid | Cold treatment | No. of egg cells | No. of cells developed into each growth stage |  |  |  | Plantlet |
| --- | --- | --- | --- | --- | --- | --- | --- |
|  |  |  | Two-celled | Cell mass | White cell colony | Plant line |  |
| **I** | - | 192 | 49 | 21 | 58 | 45 | 45 |
|  | + | 113 | 24 | 26 | 40 | 38 | 38 |
| **II** | - | 90 | 22 | 6 | 17 | 15 | 15 |
|  | + | 61 | 9 | 15 | 18 | 17 | 17 |
| **III** | - | 47 | 9 | 5 | 16 | 20 | 20 |
|  | + | 49 | 8 | 4 | 15 | 4 | 4 |
| **Total** | - | **329** | **80** | **32** | **91** | **80** | **80** |
|  | + | **223** | **41** | **45** | **73** | **59** | **59** |

**Table 5.** Ploidy levels of egg-derived plants regenerated from NBKS egg cells.

| NBKS<br>hybrid | Haploid<br>(n) | Diploid<br>(2n) | Tetraploid<br>(4n) | Aneuploidy |  |  |
| --- | --- | --- | --- | --- | --- | --- |
|  |  |  |  | 3n | 5n | 6n |
| I | 4 | 25 | 29 | 2 | 1 | 3 |
| II | 6 | 17 | 7 | 0 | 0 | 0 |
| III | 0 | 9 | 18 | 0 | 0 | 0 |
| Total | 10 | 51 | 54 | 2 | 1 | 3 |

### Genome composition of NBKS egg cell-derived rice plant

Among the NBKS egg-derived plants, one haploid plant from NBKS hybrid I egg cell, and one diploid plant and one tetraploid plant from the same NBKS hybrid II egg cell, were subjected to genome sequencing. Furthermore, genome sequencing of NB plants, KS plants, NBKS hybrids I and II, and their progenies (seedlings obtained from these hybrids) was also conducted using the information on SNPs. The origin of the sequence read was selectively identified as an NB- or KS-genome-derived read, and the ratio between NB and KS SNPs was calculated for each genome (Fig. 6). In the genomes of NBKS hybrids I and II, heterozygous profiles of SNPs from NB and KS were detected; this suggested the coexistence of alleles from both NB and KS at the same analyzed SNP-present loci because of the mixing of genetic material between the two subspecies (NB, KS, and NBKS hybrids I and II in Fig. 6). In contrast, haploid plants regenerated from an egg cell isolated from the NBKS hybrid I plant showed SNPs homologous at all the investigated loci, confirming that the haploid egg plants were derived from a single gamete after meiosis of megaspore mother cells in NBKS hybrid I (Haploid in Fig. 6).

**Figure 6.**
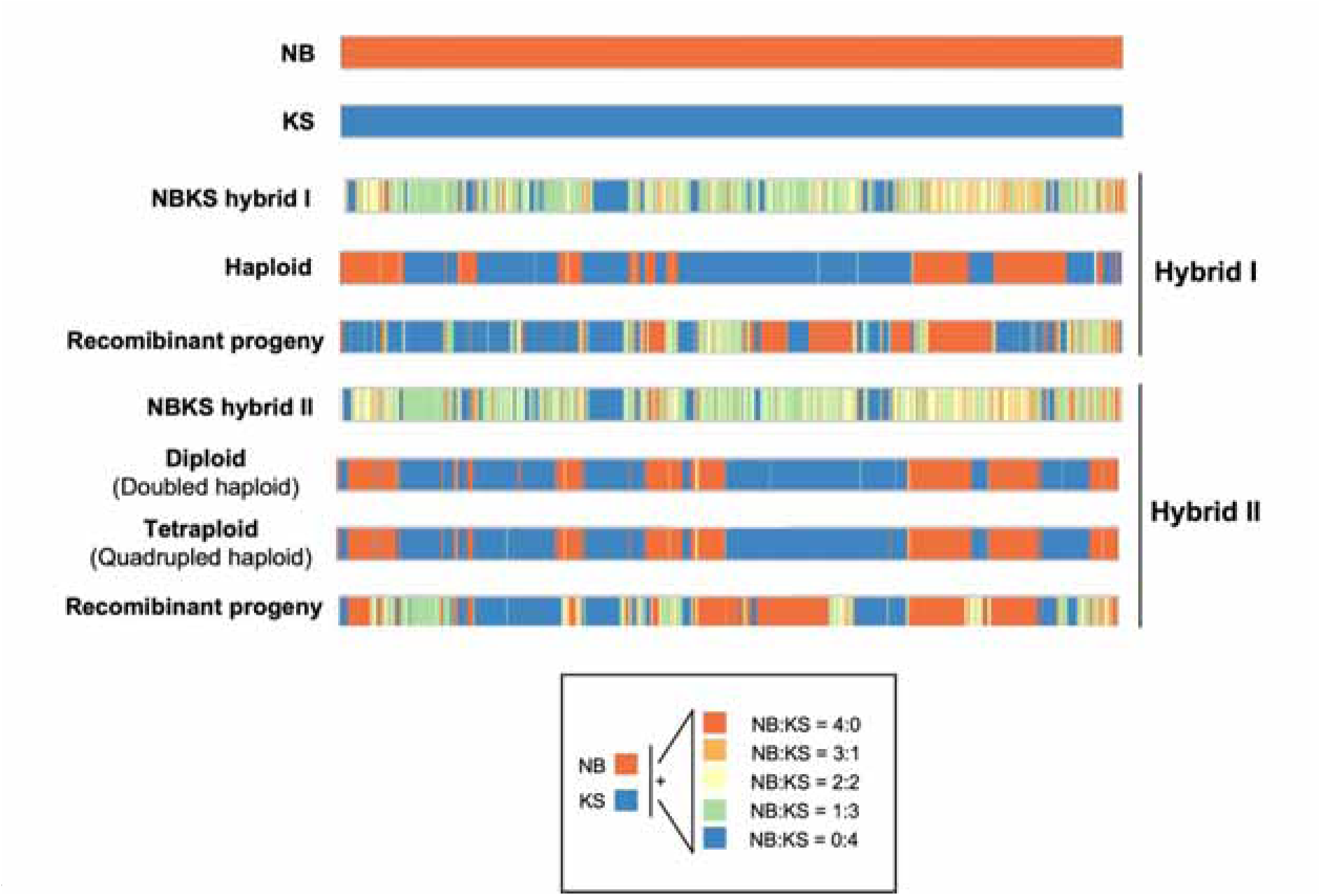
Confirmation of polyploidization of egg cell-derived plants by investigating SNPs between NB and KS in regenerated egg cells isolated from NBKS hybrid plants. Whole-genome resequencing of haploid, doubled haploid, quadrupled haploid hybrid I and II NBKS egg cell-derived plants, and recombinant progeny of the hybrid plants for detection of SNPs between NB and KS. Blue and red colors indicate the SNPs of the NB and KS alleles, respectively. Orange, yellow, and green colors represent the coexistence of both NB and KS alleles at the same SNP-containing loci.

Moreover, diploid and tetraploid plants, both originating from the same NBKS egg cell isolated from NBKS hybrid II, displayed the same homozygous patterns of SNPs throughout the genome (Diploid and Tetraploid in Fig. 6). This indicates that these plants were doubled and quadrupled haploids originating from the same egg cell, and that genome duplication occurred once during the formation of diploid NBKS egg-derived plants and twice during tetraploid plant formation from an NBKS egg cell.

### Transcriptional dynamics in egg cells post-cold treatment

Transcriptomic analysis of egg cells collected at different incubation periods after cold treatment was conducted to investigate the influence of cold stress on their gene expression profile. Briefly, isolated NB egg cells were incubated at 4 °C for 12 h or directly sampled for RNA extraction without cold treatment (C1 and C2 were biological replicates). Furthermore, the 12-h cold-treated egg cells were transferred from 4 °C and followed without (0h-1 and 0h-2) or with further incubation at 20 °C for 4 (4h-1 and 4h-2) and 12 (12h-1 and 12h-2) h.

The correlation between the transcriptomic profiles and incubation period post-cold treatment was identified via principal component analysis (Fig. 7A), which indicated that PC1 and PC2 were sufficient to separate these cells into four different groups. C1 and C2 egg cells were clustered within the range of 0h-1 and 0h-2 egg cells, suggesting that the gene expression profile of egg cells is not largely affected by cold treatment itself. The transcriptomic data of egg cells incubated for 0, 4, and 12 h post-cold-treatment were independently clustered within distinct areas (Fig. 7A). This suggests that gene expression profiles in egg cells gradually change after release from the cold treatment.

**Figure 7.**
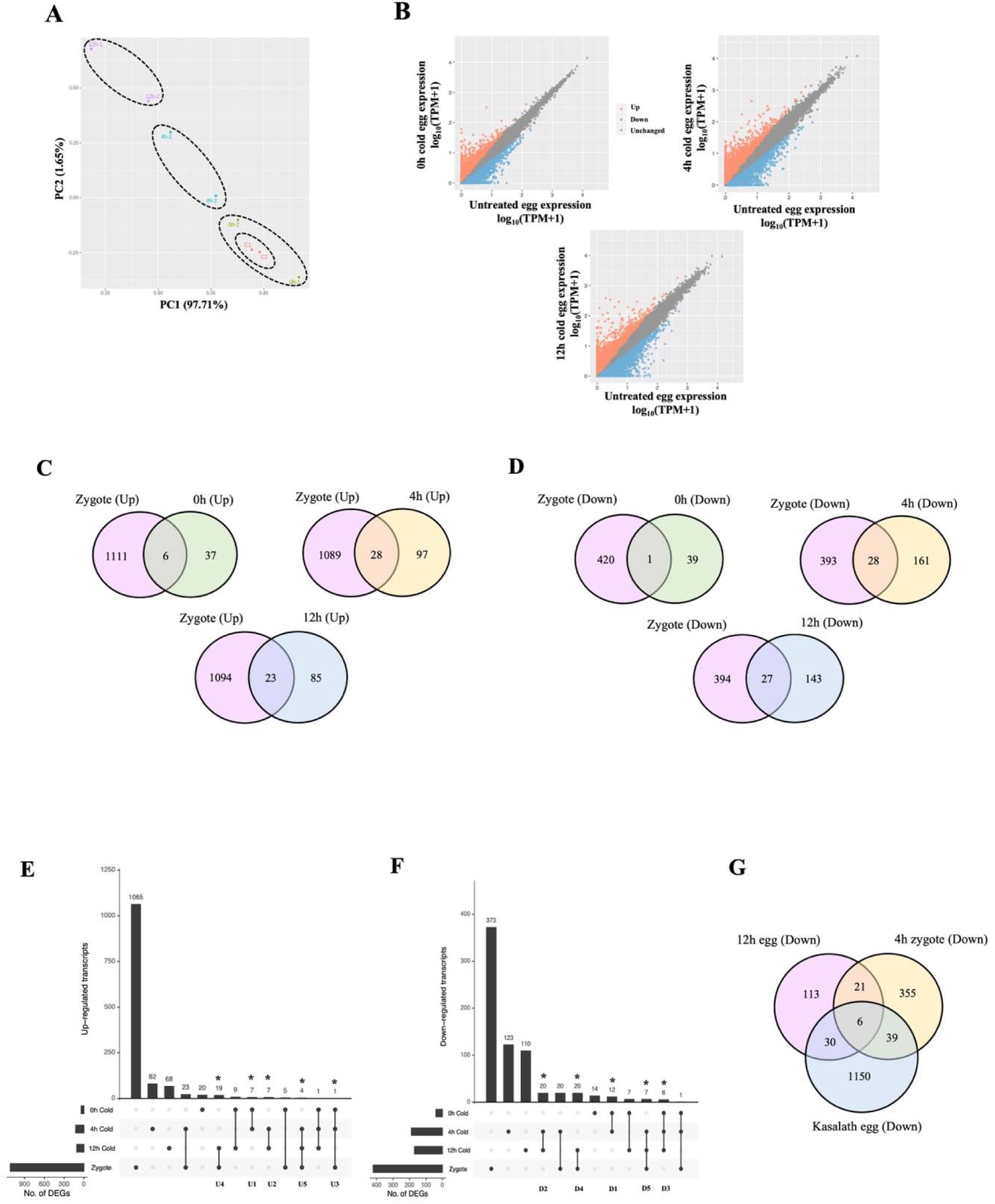
Transcriptome analysis in cold-treated egg cells. (**A**) Principal component analysis of cold-treated egg cells at different post-cold treatment incubation periods of 0, 4, and 12 h. (**B**) Scatter plots showing upregulated (red dots) and downregulated (blue dots) DEGs in each post-cold treatment incubation period at 0, 4, and 12 h compared with the untreated egg cells. (**C**) Venn diagrams indicate the commonly upregulated DEGs in the cold-treated egg cells at 0, 4, and 12 h treatments and untreated egg cells and 4h zygotes and egg cells. (**D**) Venn diagrams indicate the commonly downregulated DEGs in the cold-treated egg cells at 0, 4, and 12 h treatments and untreated egg cells and 4h zygotes and egg cells. (**E**, **F**) Set visualization diagrams for overlapping DEGs identified as upregulated and downregulated genes in cold-treated egg cells at 0h, 4h, and 12h and 4h zygotes compared to the untreated egg cells. (**E**) Transcript sets with asterisks are identified as being continuously upregulated genes at 0–4 h, 4–12 h, and 0–12 h post-cold treatment periods, 12h cold-treated eggs and 4h zygote, and 4–12h cold-treated egg and 4h zygote were extracted and classified into five categories: U1–U5, respectively. (**F**) Transcript sets with asterisks were identified as being continuously downregulated genes at 0–4h, 4–12h, and 0–12h post-cold treatment periods, 12h cold-treated eggs and 4h zygote, and 4–12h cold-treated egg and 4h zygote were extracted and classified into five categories: D1–D5, respectively. (**G**) Venn diagram indicating commonly downregulated DEGs in 12h cold-treated egg cells, 4h zygotes, and KS egg cells.

Pairwise comparisons between untreated and cold-treated egg cells were conducted to identify DEGs, including upregulated and downregulated genes, induced by cold treatment. First, we created scatter plots showing upregulated (red dots) and downregulated (blue dots) DEGs in each post-cold treatment incubation period at 0, 4, and 12 h compared with the untreated egg cells (Fig. 7B). We observed significant upregulation of 43 genes (p < 0.05) in egg cells without post-cold treatment incubation (0h eggs), 131 genes in egg cells after 4 h (4h eggs), and 110 genes in egg cells after 12 h of post-cold treatment incubation (12h eggs) compared with untreated egg cells (Fig. 7B and Supplementary Table 3). According to Gene Ontology (GO) enrichment analysis, 43 upregulated genes in 0h eggs were enriched in organic and amino acid catabolism terms (Supplementary Table 1). Studies have reported that the catabolic pathways of amino acids are activated during and after exposure to abiotic stresses, as amino acids can be used as precursors for the biosynthesis of secondary metabolites and signaling molecules that mediate stress response and tolerance to allow stress recovery and growth resumption (Batista-Silva et al., 2019; Pires et al., 2016). The highly enriched terms of upregulated genes in 4h eggs were related to chromatin remodeling, DNA packaging, amino acid metabolism, and heavy metal response (Supplementary Table 1). In addition, terms related to carbohydrate and nucleotide metabolism, heat stress response, and protein folding mechanism were enriched in 12h eggs (Supplementary Table 1), indicating that cold-treated egg cells might attempt to alleviate stress-induced molecular damage in proteins and DNA by activating protein folding mechanisms and nucleic acid metabolism.

We hypothesized that in addition to cold stress-responsive roles, these upregulated genes might also induce egg cells to undergo fertilization-independent embryogenesis. Therefore, to identify development-promoting genes among the upregulated DEGs, a comparative analysis was performed between upregulated genes in cold-treated egg cells and fertilization-induced genes in zygotes at four hours after gamete fusion (4h zygotes) (Rahman et al., 2019). We found that six genes were commonly upregulated in both 0h cold-treated egg cells and 4h zygotes (Fig. 7C). GO enrichment analysis showed that five possible stress-responsive, two protein-binding, and two transferase activity-related terms were enriched in both 0h cold-treated egg cells and zygotes (Supplementary Table 2). These results suggest that the early phase of the cellular response to cold treatment and fertilization requires a similar stress response mechanism, as a zygote also encounters developmental stress through drastic changes in metabolism and physiology after fertilization. Further, 40 genes that were commonly upregulated in 4h eggs and 4h zygotes were subjected to GO enrichment analysis (Fig. 7C). Although the enrichment of GO terms remained related to stress-responsive terms in 4h cold-treated egg cells and 4h zygotes, the enriched terms gradually shifted to amino acid metabolism, antioxidant activity, and protein-folding mechanism terms. Interestingly, in 12h cold-treated egg cells and 4h zygotes, among 24 commonly upregulated DEGs, the highly enriched terms were related to epigenetic regulation of chromatin states and protein-folding mechanisms (Fig. 7C and Supplementary Table 2), suggesting that expression of development-related genes might also be epigenetically activated after cold exposure. It was suggested that autonomous de-repression of chromatin needs to be activated to initiate a transcriptional reprogramming of egg cells to advance parthenogenesis (reviewed in Vijverberg, Ozias-Akins, and Schranz 2019). Significant enrichment of genes related to protein-folding mechanisms probably indicates the reorganization of the proteome and surveillance of proteins that are misfolded or newly synthesized because of physiological alteration of egg cells after cold treatment as well as during the conversion of an egg cell into a zygote.

Interestingly, similar to 4h zygotes, we detected an upregulation of the *OsBBML1* gene in 12h eggs compared with the control, 0h eggs, and 4h eggs, suggesting that autonomous cell division and development of egg cells exposed to cold stress might be partially induced by the upregulation of the *OsBBML1* gene (Fig. 7C and Supplementary Table 5). Since the *OsBBML1* expression level in 4h zygotes was significantly higher than that in 12h cold-treated egg cells (Fig. 7C and Supplementary Table 5), it can be concluded that some 12h cold-treated egg cells might gradually undergo an initial stage of embryogenesis, while 4h zygotes may advance towards a more progressive embryogenic stage at which reprogramming of transcriptional activity conferring embryonic development occurs, and that *de novo OsBBML1* gene expression might play a crucial role in this process.

To identify the upregulated and downregulated DEGs in cold-treated egg cells compared with the untreated cells based on the duration of post-cold treatment incubation, we created set visualization diagrams that exhibited the transcript sets of continuously upregulated or downregulated DEGs and assigned them as sets of cold treatment-induced and suppressed genes, respectively (Fig. 7E, F). For upregulated genes, all up-regulated DEGs were categorized into 13 groups based on the post-cold treatment incubation period of cold-treated egg cells and 4h zygotes at which the transcripts were upregulated DEGs compared with egg cells (Fig. 7E and Supplementary Table 3). Among the 13 groups, we selected five transcript sets that were continuously upregulated in the cold-treated egg cells at the incubation period of 0–4h, 4–12h, 0–12h, 12h and 4h zygote, and 4–12h and 4h zygote, which were classified into the upregulated gene set categories 1–5 (U1–U5), respectively (Fig. 7E and Supplementary Table 3). We speculated that the overlapping upregulated genes between cold-treated egg cells and zygotes played important roles in inducing autonomous development of unfertilized egg cells upon induction by cold treatment, as shown by the categories U4–U5 (Fig. 7E and Supplementary Table 3). Notably, in group U4, we detected an upregulation of the gene encoding the parthenogenesis-related transcription factor AP-type ERF34 (*Os04g0550200*) in 12h cold-treated egg cells and 4h zygotes, indicating the possible role of the *Os04g0550200* gene in promoting both autonomous and zygotic embryogenesis (Fig. 7E and Supplementary Table 3). However, the expression levels of this gene were found to be significantly higher in 12h eggs than in 4h zygotes (Supplementary Table 3), suggesting that autonomous development of cold-treated egg cells and zygotes might involve different pathways to initiate parthenogenetic and zygotic embryogenesis, respectively. For group U5, the *Os09g0511600* gene, which encodes the glycoside hydrolase family 1 protein, was continuously upregulated in 4h eggs, 12h eggs, and zygotes (Fig. 7E and Supplementary Table 3). Microarray data from the RiceXPro database and previous studies have reported that glycoside hydrolase enzymes, which catalyze the hydrolysis of glycosidic bonds, are required for the modification and remodeling of cell walls and are preferentially expressed in growing embryos and actively developing tissues(Sharma et al., 2013). Therefore, the upregulation of glycoside hydrolase-coding genes suggests that early acquisition of developmental competence in cold-treated egg cells and zygotes involves cell wall rearrangement, which is a fundamental factor for plant growth and development.

In addition, 12 groups of downregulated DEGs, five transcript sets that were continuously downregulated in cold-treated egg cells at the incubation period of 0–4h, 4–12h, 0–12h, 12h and 4h zygote, and 4–12h and 4h zygote, were also assigned into five categories 1–5 (D1–D5), respectively (Fig. 7E and Supplementary Table 4). We hypothesized that the overlapping downregulated genes between cold-treated egg cells and zygotes potentially suppressed fertilization-independent development and subsequent precocious embryogenesis in egg cells. Interestingly, based on the average TPM values, *Fertilization Barrier* (*FEB*) *1* and *2* genes showed a progressive decline in expression levels in cold-treated egg cells; additionally, downregulation of *FEB1* and *2* expression levels was also detected in 4h zygotes (Supplementary Table 5) (Anderson et al., 2013). This indicates that dormancy of egg cells is potentially broken upon exposure to cold stress as well as the entry of sperm nucleus, and that the pre-fertilization barrier is abolished and embryogenesis is initiated.

Moreover, we detected six commonly downregulated genes among 12h cold-treated eggs, 4h zygotes, and parthenogenetically active KS egg cells compared with NB egg cells (Fig. 7G). Notably, downregulation of *Os11t0671000*, a dormancy-associated protein, was observed in these cell types, suggesting that cold treatment might liberate the quiescent state of egg cells, thereby triggering parthenogenetic development. Downregulation of *Os05g0584200*, which encodes late embryogenesis abundant (LEA) protein, might imply that this protein is not required for early embryogenesis in cold-treated egg cells. This suggests that low LEA levels in egg cells after cold treatment allow egg cells to gain the ability to initiate early embryonic development via parthenogenesis.

## Discussion

Fertilization-independent development of female and male gametophytes has previously been reported to be effectively induced by low-temperature treatment. In this study, we demonstrated the induction of parthenogenetic development of egg cells isolated from non-parthenogenetic *japonica* rice.

Cold stress exposure is known to be an effective trigger for in vitro parthenogenesis in the sexual line of a crop plant. Upon exposure to environmental stresses, such as cold stress, sessile organisms such as plants undergo acclimation to become stress-tolerant, which allows them to survive, grow, and reproduce through alternative pathways. In several angiosperm species, the modes of reproduction can be facultative according to environmental conditions. Switching of the reproductive mode was reported in the alpine plant *Ranunculus kuepferi* upon exposure to cold stress, as it was observed that cold-treated sexual diploid plants produced significantly higher numbers of apomictic seeds than warm-treated diploids (Klatt et al., 2018). This indicates that cold stress induces a change of sexual reproduction into the reproduction mode by which the formation of asexual seeds via apomixis occurs. In the present study, we detected the incidence of autonomous cell division and development induced by cold treatment in egg cells isolated from sexual NB rice plants (Fig. 1 and 2). It is suggested that spontaneous mitosis after cold treatment in sexual haploid NB egg cells might be a mechanism to maintain reproductive fitness via parthenogenesis to acclimate to cold stress, in which both mitosis and meiosis are disrupted (Chen et al., 2011; Thakur et al., 2010). In addition to stress acclimation, parthenogenetic development is initiated by the expression of parthenogenesis-promoting genes; herein, we found an upregulation of *OsBBML1* in egg cells treated with cold stress (Fig. 7B and Supplementary Table 5). This indicates the synergistic effects of cytological and molecular responses to cold stress in activating adaptation under environmental changes by enhancing growth and reproductive ability.

We also observed the degeneration of egg cells cultured without preceding cold treatment (Fig. 1), which showed that egg cells cannot develop without the presence of proper stimuli and signals, such as the entry of a male nucleus or stress conditions. Autonomous development of cold-treated egg cells suggests that cold stress can act as a sole trigger to promote fertilization-independent embryogenesis from the isolated NB egg cells, as it was previously stated that egg cells store mRNA and proteins to prepare for embryonic development upon fertilization (Brower et al., 1981; Medvedev et al., 2011). Therefore, by employing the stored molecules, it appears that the presence of a single trigger, such as environmental stress, is sufficient to initiate embryogenesis of egg cells without the involvement of sperm cells. In actively proliferating cells, cold stress exposure has detrimental effects on cell division through spindle microtubule disorganization, and suppression, and skipping of mitosis (Moh and Alán, 1964; Brinkley and Cartwright, 1975; Lazareva et al., 2008). Therefore, cold-exposed plant cells take this opportunity to replicate their genome via endopolyploidy as an adaptation mechanism to initiate cold stress tolerance, as the resultant polyploid cells can expand in size and later proceed the cell cycle more effectively under stressful conditions (Scholes and Paige, 2015). However, in the root apical meristem of maize, the quiescent center is mitotically inactive and exists as a stem cell niche in a long-lasting halted state of cell division at the G1 phase of the cell cycle (Kerk and Feldman, 1995). Upon acclimation to cold stress, the quiescent center is triggered to break its dormancy and divide to replace the damaged cells to recover from the cold stress (Barlow and Rathfelder, 1985; Clowes and Stewart, 1967). We previously reported that rice egg cells are arrested at the G1 phase (Sukawa and Okamoto, 2018), and the current study reported that cold treatment might act as a factor that releases the cell cycle arrest in the egg cells, thereby progressing autonomous cell division without fertilization (Fig. 1).

Control of gene expression in response to cold stress can be epigenetically modulated through alteration of chromatin structure via histone acetylation, methylation, and phosphorylation (Yuan et al., 2013; Kim et al., 2015). Post-translational modification of H3 and H4 histones via acetylation of lysine residues tends to activate gene expression (Kuo et al., 1996; Zhang et al., 1999; Shahbazian and Grunstein, 2007). Cold treatment was found to trigger de-repression of chromatin via histone acetylation, which mediates the transition of chromatin from repressive to active state through *HOS15*-mediated degradation of histone deacetylase 2C (Lim et al., 2020). The subsequent hyperacetylation of histones allows chromatin to become more permissive and triggers the expression of cold-responsive *COR* genes (Park et al., 2018). Consistent with a previous study, our GO analysis showed that the highly represented group of upregulated genes in cold-treated egg cells at 4 and 12 h post-treatment were the genes encoding histone variants related to DNA packaging and nucleosome organization (Supplementary Table 1). Therefore, exposure to cold temperatures may trigger the upregulation of genes mediating chromatin reorganization in egg cells as a possible preparatory mechanism prior to cell division and development. One of the cytological features of egg cells is that their chromatin is highly condensed and repressive, as it possesses a large number of repressive histone marks, like H3K9Me2 (Fang et al., 2021). Our GO analysis results revealed a common enrichment of genes involved in epigenetic regulation in both cold-treated egg cells and 4h zygotes (Supplementary Table 1). Cold stress acclimation via chromatin structure modification mediated by epigenetic mechanisms might occur during post-treatment adaptation of cold-treated egg cells, leading to de-repression and activation of chromatin containing the genes that might be involved in promoting parthenogenesis.

Parthenogenesis is generally coupled with the restoration of the diploid number of chromosomes to affirm the viability and fertility of parthenogenetic individuals (Koltunow and Grossniklaus, 2003). Upon exposure to environmental stresses, somatic tissues in various angiosperms acquire endopolyploidy by undergoing endoreduplication without mitosis, resulting in cells with higher ploidy levels and larger sizes (Kondorosi et al., 2000; Scholes and Paige, 2015). Endopolyploidy is activated to promote the survival of plants in response to cold and environmental stresses by increasing the growth rate and metabolite storage capacity in proliferative tissues (Gegas et al., 2014; Klatt et al., 2018; Pacey et al., 2020). Notably, the results of our nuclear DNA assessment indicated that egg cells treated with cold temperature showed genomic DNA duplication, causing the transition of parthenogenetically proliferative egg cells from the haploid to diploid state at the early stage of fertilization-independent egg development (Fig. 4). A possible explanation for this is that cold stress might activate the switching of reproductive mode in egg cells from sexuality to parthenogenesis (Klatt et al., 2018). Therefore, the parthenogenetic egg cells undergo endopolyploidy to improve the survival of the subsequent egg-derived plants, as diploid and tetraploid egg plants were larger and more fertile than haploid egg plants (Fig. 3H–J, Supplementary Fig. 2 and Table 2).

Upon cold stress-induced parthenogenetic development, egg cells developed into white cell colonies and then calli with multiple green shoots, which were later regenerated into plantlets (Fig. 2). Interestingly, we often detected a variation in ploidy levels in plants regenerated from the same calli, and the number of resultant diploid and tetraploid plants was almost 1:1 (Table 2). This indicates that endopolyploidy occurs once again during callus proliferation and regeneration, and that the transition from diploid to tetraploid state begins at the late stage of parthenogenetic development of egg cells (Fig. 2 and Table 2). A previous study revealed that the induction of shoot and root regeneration in the presence of phytohormones, auxin and cytokinin, is known to cause endoreduplication and increased ploidy levels in the regenerates (Liscum and Hangarter, 1991). In addition, there is direct evidence of ploidy dynamics resulting from endopolyploidy in regenerating calli cultured on solid media that convert diploid cells into tetraploid cells (Murashige and Nakano, 1967; Torrey, 1967). However, the studies were conducted with cultivation of calli over a period of years, while the regeneration of calli into plantlets in our system only required days (Fig. 2). Therefore, it remains inconclusive whether the purpose of genome duplication via endopolyploidy in proliferating calli was to improve fitness in parthenogenetic individuals or to tolerate culture stresses. Notably, the same homozygous patterns of SNPs throughout the genome of egg-derived diploid and tetraploid plants indicated that these plants were doubled and quadrupled haploid because of endopolyploidy (Fig. 6). As weaker and infertile phenotypes were observed in haploid plants regenerated from egg cells (Supplementary Fig. 2), it can be suggested that endopolyploidy is required in cold-treated egg cells to switch the reproduction mode from sexuality to parthenogenesis to enhance viability and fertility during in vitro cultivation.

It has been previously indicated that the production of unreduced egg cells by apomeiosis and parthenogenesis is normally co-segregated(Asker and Jerling, 1992). However, this study revealed that genome duplication and parthenogenetic development are independent of each other, and these events do not occur simultaneously in in vitro cultured egg cells, as we found that the endoreduplication of egg-derived individuals occurs after the induction of parthenogenesis by cold treatment.

Previous reports regarding parthenogenesis in flowering plants have always involved apomictic seed formation, whereby embryos are derived from unreduced egg cells resulting from apomeiosis, which produces genetically identical offspring (Koltunow and Grossniklaus, 2003). In this study, we provide new insights into the production of plants directly from haploid egg cells derived from meiosis without apomeiotic events; hence, the plants regenerated from these egg cells are genetically variable from the mother because of genetic recombination during meiosis. Therefore, egg-derived progeny with variable traits can be obtained without genetic contributions from the male parent, and offspring with desired traits can be obtained through further selection. Furthermore, as cold treatment is known to be beneficial for fertilization-independent embryogenesis derived from reproductive structures (Sibi et al., 2001; Gürel et al., 2000), exposure to low temperatures prior to in vitro culture of isolated egg cells may be a potential technique to physically induce the generation of desired crop plants from unfertilized gametes.

## Materials and Methods

### Plant materials and cultures of the isolated egg cells exposed to cold temperature

*Oryza sativa* L. cv. Nipponbare (NB), *O. sativa* L. cv. Kasalath (KS) and NB-KS hybrid rice plants were grown in an environmental chamber (K30-7248; Koito Industries, Yokohama, Japan) at 26 °C under a 13 h light/11 h dark photoperiod. To visualize the nuclei in egg cells and multicellular structures, transformed KS rice plants expressing H2B-GFP were prepared as previously described (Abiko et al., 2013). Egg cells were isolated from rice flowers as previously described (Uchiumi et al., 2006, 2007). To investigate cold treatment-induced parthenogenesis, isolated egg cells were incubated with or without cold treatment at 4 °C for 12 h and subsequently cultured in N6Z medium.

To create NB-KS hybrid plants, an NB egg cell was fertilized with a KS sperm cell using electrofusion, as described previously (Uchiumi et al., 2007). The resultant NB-KS hybrid zygotes were cultured into plantlets and termed NBKS hybrids. Egg cells were isolated from the NBKS hybrid and subsequently incubated with or without cold pretreatment (4 °C for 12 h). The incubated egg cells were then cultured in the N6Z medium.

### Microscopic observation

Cellular features and developmental profiles of the egg cells and parthenogenetic multicellular structures were observed using a BX-71 inverted microscope (Olympus, Tokyo, Japan). Digital images of egg cells and the resultant multicellular structures were obtained using a cooled charge-coupled device camera (Penguin 600CL; Pixcera, CA, USA) and the InStudio software (Pixcera). In addition to the BX-71 inverted fluorescence microscope, egg cells and multicellular structures expressing green fluorescent protein (GFP) fusion proteins or stained with fluorescent probes were observed using an LSM 710 CLS microscope (Carl Zeiss, Jena, Germany), with excitation and emission wavelengths specific to each type of fluorophores as described below. The intracellular fluorescent signal of H2B-GFP proteins was observed under a BX-71 inverted fluorescence microscope (Olympus) at 460–490-nm excitation and 510–550-nm emission wavelengths (U-MWIBA2 mirror unit; Olympus). To quantify the relative DNA amount in the cells of parthenogenetic multicellular structure using mDAPI (Sukawa and Okamoto, 2018), the fluorescent signal of the nuclei of egg cells and multicellular structures at 2-celled, 4- to 8-celled, and 10- to 20-celled stages were stained with mDAPI and observed using the BX-71 inverted microscope (Olympus) with 360–370-nm excitation and 420–460-nm emission wavelengths (U-MNUA mirror unit; Olympus).

### Quantification of the relative amount of nuclear DNA in egg cells and parthenogenetically developed multicellular structures

The isolated NB egg cells and multicellular structures at 2-celled, 4- to 8-celled, and 10- to 20-celled stages were subjected to DNA staining using the mDAPI method as previously described (Sukawa and Okamoto, 2018). Briefly, the egg cells and multicellular structures were placed in staining droplets containing a mixture of nuclear extraction buffer and DAPI at a ratio of 1:5 and incubated for 2 min. The cells were then observed under a BX-71 inverted microscope (Olympus) and the fluorescence intensity was quantified using ImageJ software.

### Flow cytometry analysis

The nuclear DNA content (ploidy level) of plants regenerated from cold-treated egg cells was measured with flow cytometry using CyFlow CCA (Partec, Muenster, Germany) and a QuantumStain NA UV2 Kit (Quantum Analysis, Muenster, Germany). Fresh leaves (22–25 mm^2^) were chopped with a sharp razor in 100 ll of the kit solution, and 100 ll more was added after chopping. The samples were incubated for 5 min, and the crushed tissue suspension was filtered through a 30 lm nylon mesh (Partec). The samples were subsequently loaded into a ploidy analyzer. Leaf samples from diploid wild-type NB plants (2n = 24, *Oryza sativa* L. cv NB) were used as the internal controls. Fluorescence intensities were measured and plotted in real-time. The peak and mean values were calculated using the FloMax software.

### Preparation of genomic DNA for whole-genome resequencing

Leaf samples derived from NB, KS, NBKS hybrids I and II, NBKS hybrids I and II recombinant progeny, NBKS hybrid I egg-derived haploid, and NBKS hybrid II egg-derived diploid and tetraploid plants were collected. Approximately 200 mg of leaf material was ground in liquid nitrogen, and the resultant powder was transferred and redissolved in lysis buffer provided by the Nucleospin Plant II kit (Macherey-Nagel, Clonetech, CA, USA). Genome isolation was performed according to the manufacturer’s instructions. After eluting the purified genomic DNA, its quality and quantity were verified using a Nanodrop 2000. Genomic library construction was performed using the Nextera DNA Flex Library Prep kit (Illumina, San Diego, CA, USA) according to the manufacturer’s instructions. Thereafter, the obtained genomic libraries were purified with Agencourt AMPure XP and subjected to quantity and quality determination using a Qubit 3 Fluorometer with a Qubit dsDNA HS Assay Kit (Thermo Fisher Scientific, Inc., Waltham, MA, USA) and an Agilent 2100 BioAnalyzer with a high-sensitivity DNA chip (Agilent Technologies, Santa Clara, CA, USA). After checking the quality and quantity of the purified libraries, 150-bp paired-end reads were generated on an Illumina HiSeq platform (Illumina) with an average depth of approximately 20-fold coverage for each sample at Macrogen-Japan (Tokyo, Japan).

### Calculation of SNP counts in genomic sequences from the KS and NB genomes

The quality of Illumina reads was evaluated using FastQC (v0.11.8) (Simon). Preprocessing of the reads included removing the adapter and low-quality sequences was conducted using Cutadapt (v2.10) (Martin, 2011). The remaining reads were mapped to the NB genome sequences (Os-Nipponbare-Reference-IRGSP-1.0) available at the Rice Annotation Project Database (RAP-DB) (Sakai et al., 2013; Kawahara et al., 2013) using BWA (v0.7.17) (Li and Durbin, 2009). PCR duplications were marked with Picard Toolkit (https://broadinstitute.github.io/picard/), then variant calling was performed by GATK (v4.2.2.0) (Auwera and O’Connor, 2020). The detected variants were filtered using VCFtools (v0.1.16) (Danecek et al., 2011) with the following options: ‘—minDP 10’ and ‘—max-missing 1’. Genotyping was performed using an R package, UPDOG (v2.0.2) (Gerard et al., 2018) and high-confidence SNPs were selected (prop_mis < 0.05).

### Preparation of lysates from cold-treated egg cells for mRNA sequencing

Isolated NB egg cells were incubated at 4 °C for 12 h or directly sampled for RNA extraction without cold treatment. Subsequently, the 12-h cold-treated egg cells were transferred from 4 °C and followed without or with further incubation at 20 °C for 4 and 12 h. Two independent biological replicates of 8–10 egg cells derived from each condition were subjected to RNA extraction. Isolated egg cells with or without cold treatment were transferred to mannitol droplets, and adjusted the osmolarity to 370 mOsmol kg^−1^ H_2_O on coverslips. The egg cells were washed three times by transferring them to fresh droplets of mannitol solution. After washing, each egg cell sample was transferred to a lysis buffer supplied in the SMART-Seq HT kit (Takara Bio, Shiga, Japan). The lysates were immediately used for cDNA synthesis or frozen in liquid nitrogen and stored at -80 °C. Thereafter, cDNA was synthesized and amplified, and libraries were constructed.

### cDNA synthesis and library preparation

cDNA preparation and library construction were performed as previously described (Deushi et al., 2021). Briefly, cDNA was synthesized and amplified from cell lysates using a SMART-Seq HT Kit (Takara Bio) according to the manufacturer’s instructions. The resulting amplified cDNA was purified using Agencourt AMPure XP (Beckman Coulter, Brea, CA, USA). The quality and quantity of the purified cDNA were determined using a Qubit 3 Fluorometer with a Qubit dsDNA HS Assay Kit (Thermo Fisher Scientific) and an Agilent 2100 BioAnalyzer with a high-sensitivity DNA chip (Agilent Technologies). Sequencing libraries were prepared from the amplified cDNA using the Nextera XT DNA Library Prep Kit (Illumina) according to the SMART-Seq HT Kit instruction manual, after which they were purified using Agencourt AMPure XP. After checking the quality and quantity of the purified libraries using the above-mentioned procedures for the purified cDNA, the prepared libraries were sequenced on an Illumina HiSeq platform (Illumina) at Macrogen-Japan (Tokyo, Japan) to produce 150-bp paired-end reads.

### Analysis of transcriptome data

The quality of Illumina reads was evaluated using FastQC (v0.11.8) (Simon). Preprocessing of the reads included removing the adapter, poly-A, and low-quality sequences was conducted using Cutadapt (v2.10) (Martin, 2011). The remaining high-quality reads were mapped to the NB transcript sequences (version IRGSP-1.0) available at the Rice Annotation Project Database (RAP-DB) (Sakai et al., 2013; Kawahara et al., 2013), and the TPM was calculated using RSEM (v1.3.1) (Li and Dewey, 2011) with Bowtie2 (v 2.3.5.1) (Langmead and Salzberg, 2012). The differentially expressed genes (DEGs) between the untreated and cold-treated egg cells derived from different post-cold treatment incubation periods were detected using an R package, TCC (Sun et al., 2013). Genes with a false discovery rate (FDR; q-value) <0.05 were extracted as DEGs. Set visualization of the overlapping DEGs based on the time of 20 °C-incubation after cold treatment and zygotes(Rahman et al., 2019) was generated using the UpsetR package (Conway et al., 2017), part of the R software. The data were statistically analyzed using a hypergeometric test in ShinyGO v0.66 (Ge, 2020).

## Acknowledgements

We thank Ms. T. Mochizuki (Tokyo Metropolitan University) for isolating rice egg cells, Mr. D. Akasaka for providing transgenic rice plants (cv. Kasalath) (Tokyo Metropolitan University) and the RIKEN Bio Resource Center (Tsukuba, Japan) for providing cultured rice cells (Oc line). We would also like to thank Dr. A. Kinoshita and Dr. E. Toda for providing valuable advice on R-software operation. Computations were partially performed on the NIG supercomputer at the ROIS National Institute of Genetics.

## Author contributions

K.R. K.T. and T.O. designed the experiments; K.R. performed most of the experiments; K.T. performed nuclear DNA quantification using mDAPI staining and cold-treated egg cell developmental profile monitoring, respectively. S.K. and K.Y. performed analyses of SNPs and transcriptome data; K.R. and T.O. conceived the project and wrote the article.

## References

Abiko, M., Maeda, H., Tamura, K., Hara-Nishimura, I., and Okamoto, T. (2013). Gene expression profles in rice gametes and zygotes: Identification of gamete-enriched genes and up-or down-regulated genes in zygotes after fertilization. J. Exp. Bot. 64: 1927–1940.

Anderson, S.N., Johnson, C.S., Jones, D.S., Conrad, L.J., Gou, X., Russell, S.D., and Sundaresan, V. (2013). Transcriptomes of isolated Oryza sativa gametes characterized by deep sequencing: Evidence for distinct sex-dependent chromatin and epigenetic states before fertilization. Plant J. 76: 729–741.

Asker, S.E. and Jerling, L. (1992). Apomixis in Plants 1st ed. (CRC press: Boca Raton, FL).

Auwera, G.A. Van der and O’Connor, B.D. (2020). Genomics in the Cloud (O’Reilly Media, Inc.).

Avise, J.C. (2008). Clonality: The Genetics, Ecology and Evolution of Sexual Abstinence in Vertebrate Animals (Oxford University Press: Oxford).

Barlow, P.W. and Rathfelder, E.L. (1985). Cell division and regeneration in primary root meristems of Zea mays recovering from cold treatment. Environ. Exp. Bot. 25: 303–314.

Baroux, C. and Grossniklaus, U. (2015). Chapter Ten - The Maternal-to-Zygotic Transition in Flowering Plants: Evidence, Mechanisms, and Plasticity. Curr. Top. Dev. Biol. 113: 351–371.

Batista-Silva, W., Heinemann, B., Rugen, N., Nunes-Nesi, A., Araújo, W.L., Braun, H.P., and Hildebrandt, T.M. (2019). The role of amino acid metabolism during abiotic stress release. Plant Cell Environ. 42: 1630–1644.

Brinkley, B.R. and Cartwright, J. (1975). Cold-Labile and Cold-Stable Microtubules in the Mitotic Spindle of Mammalian Cells. Ann. N. Y. Acad. Sci. 253: 428–439.

Brower, P.T., Gizang, E., Boreen, S.M., and Schultz, R.M. (1981). Biochemical studies of mammalian oogenesis: synthesis and stability of various classes of RNA during growth of the mouse oocyte in vitro. Dev. Biol. 86: 373–383.

Chen, J., Strieder, N., Krohn, N.G., Cyprys, P., Sprunck, S., Engelmann, J.C., and Dresselhaus, T. (2017). Zygotic genome activation occurs shortly after fertilization in maize. Plant Cell 29: 2106–2125.

Chen, N., Xu, Y., Wang, X., Du, C., Du, J., Yuan, M., Xu, Z., and Chong, K. (2011). OsRAN2, essential for mitosis, enhances cold tolerance in rice by promoting export of intranuclear tubulin and maintaining cell division under cold stress. Plant, Cell Environ. 34: 52–64.

Clowes, F.A.L. and Stewart, H.E. (1967). Recovery From Dormancy in Roots. New Phytol. 66: 115–123.

Conner, J.A., Mookkan, M., Huo, H., Chae, K., and Ozias-Akins, P. (2015). A parthenogenesis gene of apomict origin elicits embryo formation from unfertilized eggs in a sexual plant. Proc. Natl. Acad. Sci. U. S. A. 112: 11205–11210.

Conner, J.A., Podio, M., and Ozias-Akins, P. (2017). Haploid embryo production in rice and maize induced by PsASGR-BBML transgenes. Plant Reprod. 30: 41–52.

Conway, J.R., Lex, A., and Gehlenborg, N. (2017). Genome analysis UpSetR: an R package for the visualization of intersecting sets and their properties. 33: 2938–2940.

Danecek, P. et al. (2011). The variant call format and VCFtools. 27: 2156–2158.

Deushi, R., Toda, E., Koshimizu, S., Yano, K., and Okamoto, T. (2021). Effect of Paternal Genome Excess on the Developmental and. Plants: 1–13.

Fang, H., Shao, Y., and Wu, G. (2021). Reprogramming of Histone H3 Lysine Methylation During Plant Sexual Reproduction. Front. Plant Sci. 12: 1–17.

Garcia-Aguilar, M., Michaud, C., Leblanc, O., and Grimanelli, D. (2010). Inactivation of a DNA methylation pathway in maize reproductive organs results in apomixis-like phenotypes. Plant Cell 22: 3249–3267.

Ge, S.X. (2020). ShinyGO: a graphical gene-set enrichment tool for animals and plants. 36: 2628–2629.

Gegas, V.C., Wargent, J.J., Pesquet, E., Granqvist, E., Paul, N.D., and Doonan, J.H. (2014). Endopolyploidy as a potential alternative adaptive strategy for Arabidopsis leaf size variation in response to UV-B. J. Exp. Bot. 65: 2757–2766.

Gerard, D., Felipe, L., Ferrão, V., Augusto, A., and Garcia, F. (2018). Genotyping Polyploids from Messy Sequencing Data. 210: 789–807.

Gürel, S., Gürel, E., and Kaya, Z. (2000). Doubled haploid plant production from unpollinated ovules of sugar beet (Beta vulgaris L.). Plant Cell Rep. 19: 1155–1159.

Hand, M.L. and Koltunow, A.M.G. (2014). The genetic control of apomixis: Asexual seed formation. Genetics 197: 441–450.

Kawahara, Y. et al. (2013). Improvement of the Oryza sativa Nipponbare reference genome using next generation sequence and optical map data.: 1–10.

Kerk, N.M. and Feldman, L.J. (1995). A biochemical model for the initiation and maintenance of the quiescent center: Implications for organization of root meristems. Development 121: 2825–2833.

Khanday, I., Skinner, D., Yang, B., Mercier, R., and Sundaresan, V. (2019). A male-expressed rice embryogenic trigger redirected for asexual propagation through seeds. Nature 565: 91–95.

Kim, J.M., Sasaki, T., Ueda, M., Sako, K., and Seki, M. (2015). Chromatin changes in response to drought, salinity, heat, and cold stresses in plants. Front. Plant Sci. 6: 1–12.

Klatt, S., Schinkel, C.C.F., Kirchheimer, B., Dullinger, S., and Hörandl, E. (2018). Effects of cold treatments on fitness and mode of reproduction in the diploid and polyploid alpine plant Ranunculus kuepferi (Ranunculaceae). Ann. Bot. 121: 1287–1298.

Koltunow, A.M. and Grossniklaus, U. (2003). Apomixis: A Developmental Perspective. Annu. Rev. Plant Biol. 54: 547–574.

Kondorosi, E., Roudier, F., and Gendreau, E. (2000). Plant cell-size control: Growing by ploidy? Curr. Opin. Plant Biol. 3: 488–492.

Kumlehn, J., Kirik, V., Czihal, A., Altschmied, L., Matzk, F., Lörz, H., and Bäumlein, H. (2001). Parthenogenetic egg cells of wheat: Cellular and molecular studies. Sex. Plant Reprod. 14: 239–243.

Kuo, M.H., Brownell, J.E., Sobel, R.E., Ranalli, T.A., Cook, R.G., Edmondson, D.G., Roth, S.Y., and Allis, C.D. (1996). Transcription-linked acetylation by Gcn5p of histones H3 and H4 at specific lysines. Nature 383: 269–272.

Langmead, B. and Salzberg, S.L. (2012). Fast gapped-read alignment with Bowtie 2. 9: 357–360.

Lazareva, E.M., Chentsov, Y.S., and Smirnova, E.A. (2008). The effect of low temperature on the microtubules in root meristem cells of spring and winter cultivars of wheat Triticum aestivum L. Cell tissue biol. 2: 436–450.

Li, B. and Dewey, C.N. (2011). RSEM: accurate transcript quantification from RNA-Seq data with or without a reference genome.

Li, H. and Durbin, R. (2009). Fast and accurate short read alignment with Burrows – Wheeler transform. 25: 1754–1760.

Lim, C.J. et al. (2020). The Histone-modifying complex PWR/HOS15/HD2C epigenetically regulates cold tolerance1[OPEN]. Plant Physiol. 184: 1097–1111.

Liscum, E. and Hangarter, R.P. (1991). Manipulation of Ploidy Level in Cultured Haploid Petunia Tissue by Phytohormone Treatments. J. Plant Physiol. 138: 33–38.

Liu, X.-Q., Shi, J.-J., Fan, H., Jiao, J., Gao, L., Tan, L., Nagawa, S., and Wang, D.-Y. (2020). Nuclear DNA replicates during zygote development in Arabidopsis and *Torenia fournieri*. Plant Physiol.: 137–145.

Martin, M. (2011). Cutadapt removes adapter sequences from high-throughput sequencing reads. EMBnet 17: 10–12.

Medvedev, S., Pan, H., and Schultz, R.M. (2011). Absence of MSY2 in mouse oocytes perturbs oocyte growth and maturation, RNA stability, and the transcriptome. Biol. Reprod. 85: 575–583.

Mogensen, H.L. and Holm, P.B. (1995). Dynamics of nuclear DNA quantities during zygote development in barley. Plant Cell 7: 487–494.

Moh, C.C. and Alán, J.J. (1964). The Effect of Low Temperature on Mitosis in the Root Tips of Beans. Caryologia 17: 409–415.

Murashige, T. and Nakano, R. (1967). Chromosome Complement as a Determinant of the Morphogenic Potential of Tobacco Cells. Am. J. Bot. 54: 963.

Pacey, E.K., Maherali, H., and Husband, B.C. (2020). Endopolyploidy is associated with leaf functional traits and climate variation in Arabidopsis thaliana. Appl. Plant Sci. 107: 993–1003.

Park, J. et al. (2018). Epigenetic switch from repressive to permissive chromatin in response to cold stress. Proc. Natl. Acad. Sci. U. S. A. 115.

Pillot, M., Baroux, C., Vazquez, M.A., Autran, D., Leblanc, O., Vielle-Calzada, J.P., Grossniklaus, U., and Grimanelli, D. (2010). Embryo and endosperm inherit distinct chromatin and transcriptional states from the female gametes in Arabidopsis. Plant Cell 22: 307–320.

Pires, M. V., Pereira Júnior, A.A., Medeiros, D.B., Daloso, D.M., Pham, P.A., Barros, K.A., Engqvist, M.K.M., Florian, A., Krahnert, I., Maurino, V.G., Araújo, W.L., and Fernie, A.R. (2016). The influence of alternative pathways of respiration that utilize branched-chain amino acids following water shortage in Arabidopsis. Plant Cell Environ. 39: 1304–1319.

Pulido, A., Bakos, F., Devic, M., Barnabás, B., and Olmedilla, A. (2009). HvPG1 and ECA1: Two genes activated transcriptionally in the transition of barley microspores from the gametophytic to the embryogenic pathway. Plant Cell Rep. 28: 551–559.

Pupilli, F. and Barcaccia, G. (2012). Cloning plants by seeds: Inheritance models and candidate genes to increase fundamental knowledge for engineering apomixis in sexual crops. J. Biotechnol. 159: 291–311.

Rahman, M.H., Toda, E., Kobayashi, M., Kudo, T., Koshimizu, S., Takahara, M., Iwami, M., Watanabe, Y., Sekimoto, H., Yano, K., and Okamoto, T. (2019). Expression of genes from paternal alleles in rice zygotes and involvement of OsASGR-BBML1 in initiation of zygotic development. Plant Cell Physiol. 60: 725–737.

Sailer, C., Schmid, B., and Grossniklaus, U. (2016). Apomixis allows the transgenerational fixation of phenotypes in hybrid plants. Curr. Biol. 26: 331–337.

Sakai, H. et al. (2013). Rice Annotation Project Database (RAP-DB): An Integrative and Interactive Database for Rice Genomics Special Focus Issue – Databases. 54.

Scholes, D.R. and Paige, K.N. (2015). Plasticity in ploidy: A generalized response to stress. Trends Plant Sci. 20: 165–175.

Shahbazian, M.D. and Grunstein, M. (2007). Functions of Site-Specific histone acetylation and deacetylation. Annu. Rev. Biochem. 76: 75–100.

Sharma, R., Cao, P., Jung, K.H., Sharma, M.K., and Ronald, P.C. (2013). Construction of a rice glycoside hydrolase phylogenomic database and identification of targets for biofuel research. Front. Plant Sci. 4: 1–15.

Sibi, M.L., Kobaissi, A., and Shekafandeh, A. (2001). Green haploid plants from unpollinated ovary culture in tetraploid wheat (Triticum durum Defs.). Euphytica 122: 351–359.

Simon, A. FastQC: A Quality Control Tool for High Throughput Sequence Data.

Sukawa, Y. and Okamoto, T. (2018). Cell cycle in egg cell and its progression during zygotic development in rice. Plant Reprod. 31: 107–116.

Sun, J., Nishiyama, T., Shimizu, K., and Kadota, K. (2013). TCC: an R package for comparing tag count data with robust normalization strategies TCC: an R package for comparing tag count data with robust normalization strategies.

Thakur, P., Kumar, S., Malik, J.A., Berger, J.D., and Nayyar, H. (2010). Cold stress effects on reproductive development in grain crops: An overview. Environ. Exp. Bot. 67: 429–443.

Toki, S., Hara, N., Ono, K., Onodera, H., Tagiri, A., Oka, S., and Tanaka, H. (2006). Early infection of scutellum tissue with Agrobacterium allows high-speed transformation of rice. Plant J. 47: 969–976.

Torrey, J.G. (1967). Morphogenesis in Relation to Chromosomal Constitution in Long-term Plant Tissue Cultures. Physiol. Plant. 20: 265–275.

Tsunewaki, K. and Mukai, Y. (1990). Wheat Haploids Through the Salmon Method. 13: 460–478.

Uchiumi, T., Komatsu, S., Koshiba, T., and Okamoto, T. (2006). Isolation of gametes and central cells from *Oryza sativa* L. Sex. Plant Reprod. 19: 37–45.

Uchiumi, T., Uemura, I., and Okamoto, T. (2007). Establishment of an in vitro fertilization system in rice (*Oryza sativa* L.). Planta 226: 581–589.

Underwood, C.J. et al. (2022). A PARTHENOGENESIS allele from apomictic dandelion can induce egg cell division without fertilization in lettuce. Nat. Genet. 54: 84–93.

Vijverberg, K., Ozias-Akins, P., and Schranz, M.E. (2019). Identifying and engineering genes for parthenogenesis in plants. Front. Plant Sci. 10.

Wuest, S.E., Vijverberg, K., Schmidt, A., Weiss, M., Gheyselinck, J., Lohr, M., Wellmer, F., Rahnenführer, J., von Mering, C., and Grossniklaus, U. (2010). Arabidopsis Female Gametophyte Gene Expression Map Reveals Similarities between Plant and Animal Gametes. Curr. Biol. 20: 506–512.

Yuan, L., Liu, X., Luo, M., Yang, S., and Wu, K. (2013). Involvement of histone modifications in plant abiotic stress responses. J. Integr. Plant Biol. 55: 892–901.

Zhang, W., Bone, J.R., Edmondson, D.G., Turner, B.M., Roth, S.Y., and Annunziato, A.T. (1999). Essential and redundant functions of histone acetylation revealed by mutation of target lysines and loss of Gcn5p acetyltransferase. Chemtracts 12: 748–754.

